# BasalCell: A project scaffold generator for bioinformatics analysis

**DOI:** 10.64898/2026.05.27.720396

**Authors:** Yuji Okano, Tetsuo Ishikawa, Kazuhiro Sakurada

## Abstract

In the current bioinformatics landscape, where R-centric and Python-centric ecosystems coexist and overlap, there is an increasing demand for the organic integration of these disparate development environments. As bioinformatics practices become ubiquitous, it is crucial to lower the technical barriers for biologists to adopt software engineering standards—such as version control, environment reproducibility, code readability, continuous integration/continuous deployment (CI/CD), and comprehensive documentation—which often present significant implementation hurdles for biologists with limited programming experience.

To address these challenges, we developed `BasalCell` (https://github.com/yo-aka-gene/BasalCell), a project scaffolding system designed to provide a standardized, easily reproducible template that integrates these essential features by default—enabling researchers to seamlessly manage multi-language environments and automate rigorous development workflows, ultimately fostering greater transparency and reliability in biological data science.

## Introduction

The rapid proliferation of bioinformatics tools, particularly the rise of Python’s `scverse` [1] alongside historically dominant R, necessitates multi-language workflows in modern bioinformatics [2]. However, managing complex environments and robust version control poses a formidable barrier for biologists lacking extensive programming expertise.

To address this challenge, we developed BasalCell, a simple project scaffold generator that streamlines setting up integrative, reproducible, multi-language environments.

## Building professional development environments within a click

The core philosophy of BasalCell is to democratize standard software engineering practices for environment configuration, enabling biologists of any skill level to easily and reproducibly set up their workspaces (Figures 1a–1b). BasalCell leverages the `cookiecutter` framework [3] to reproduce professional coding environments through a few user prompts (Figure 1c). By integrating multi-language frameworks such as `Poetry` [4] and `renv` [5] within a unified `Mamba` [6] environment, the platform ensures rigorous control over language and package versions for Python and R, offering simplified initiation and reproducibility (see also Methods B.1.2 for details).

**Figure 1.**
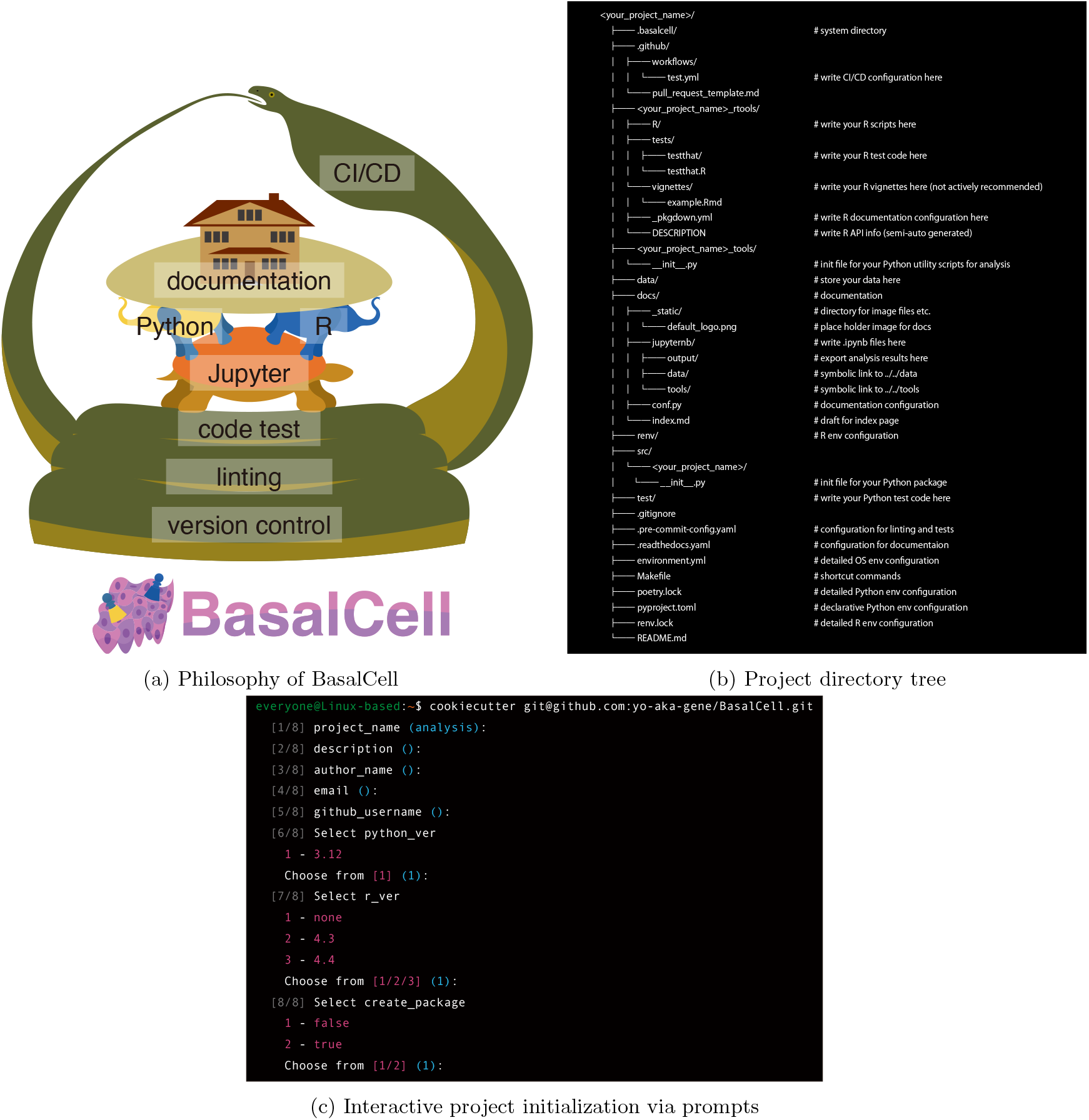
Overview of the BasalCell framework. **a**: The core philosophy of BasalCell: providing an ecosystem that harmonizes Python and R on a Jupyter foundation and seamlessly integrates them with automated documentation. This workflow is underpinned by softwaredevelopment-level environmental robustness and continuously monitored via server-side CI/CD pipelines. **b**: An illustrative example of a project directory tree generated by BasalCell (configured with options for R language usage and package development mode). For clarity, certain auxiliary files are omitted. For reference, the actual project directory generated under this configuration (named `BasalCellBlank`) is publicly deposited on GitHub (https://github.com/yo-aka-gene/BasalCellBlank). **c**: An illustrative example of the interactive CLI prompt used to generate a BasalCell project. The complete environment is scaffolded by executing a single command and answering eight intuitive prompts. Light blue braces indicate the default values for each variable. See Methods B.1.1 for detailed notes regarding Python version constraints. Note that the actual terminal appearance may vary depending on individual user configurations.

## An integrated multi-language environment

While resolving backend dependencies is crucial, providing a unified integrated development environments (IDE) is equally important for practical data analysis. Conventionally, managing disparate language-specific IDEs in standalone repositories or juxtaposing them in a pre-configured environment—an approach we previously employed [7]—complicates project synchronization and reproducibility (see also https://github.com/yo-aka-gene/algebra_ver822).

In contrast, BasalCell facilitates the entire workflow within a unified virtual environment for Python and R via a single JupyterLab-based notebook interface [8], from the initial directory preparation. Furthermore, BasalCell orchestrates multi-platform configurations via a unified, queryable dataframe with Markdown export options, enabling seamless version tracking for both automated system execution and human readability.

## Automated CI/CD on GitHub

Once a reproducible environment is established, maintaining code quality during collaborative development becomes the next critical challenge. When multiple researchers collaborate on data analysis, challenges related to code readability and CI/CD troubleshooting can become significant barriers. Since expertise in CI/CD—the automated workflows that systematically test, format, and verify code—is often outside the primary scope of biological research, it represents a steep learning curve for biologists who are new to programming.

To guarantee code quality across languages, Basal-Cell integrates standard Python and R frameworks for unit testing and linting, seamlessly orchestrated via `pre-commit` [9] hooks and GitHub Actions.

Notably, while BasalCell fully automates routine tasks like linting, it provides significant flexibility regarding test-driven development (TDD) by offering a preconfigured environment without enforcing rigid testing protocols (as detailed in Methods B.1.3). Depending on development constraints and project objectives, developers are free to adjust the target, stringency, and scope of their tests. Furthermore, the framework natively supports `doctest`, offering a lightweight and simplified alternative for quick function validation.

Given these robust TDD and CI/CD foundations suitable for production-level Python packages, BasalCell explicitly supports package development workflows. By setting the `create package` argument to `true` (Figure 1c), the framework generates a directory structure tailored for package development (e.g., initializing a `src`/ directory; see Figure 1b for details). This enables developers to seamlessly publish their code as a Python package to PyPI via Poetry.

## Making every single project documentation-ready

Beyond ensuring internal code quality through testing, communicating the analytical rationale to the broader scientific community is essential for methodological transparency. A recurring issue in bioinformatics is the poor readability of code repositories, where loosely organized raw scripts make it difficult to verify expected outputs. BasalCell addresses this by utilizing the notebook interface in JupyterLab, where outputs are displayed directly beneath the corresponding code blocks, significantly enhancing immediate visual context for reviewers.

However, even with readable notebooks, a project’s clarity can suffer if custom function scripts are disorganized. To resolve this, BasalCell adopts `Sphinx` [10] to semi-automatically generate online documentation from source code docstrings. By integrating notebooks and raw scripts into the documentation, the framework enhances the overall comprehensibility of the codebase across the entire repository.

While Sphinx is standard for documenting software packages, BasalCell extends this capability to custom data analysis modules to resolve the common issue of poor repository readability. Specifically, the `tools`/ directory is designated for utility functions and auxiliary scripts—components typically excluded from the core package but essential for methodological transparency. By automatically targeting these scripts for documentation, BasalCell ensures that auxiliary logic remains as interpretable as the primary codebase. We have previously demonstrated the efficacy of this approach even in a data analysis repository [11], which significantly improved methodological transparency (https://takemura-hgf.readthedocs.io). Althoguh we previously relied on Python wrappers to document R scripts, BasalCell now natively supports dual-language documentation by hyperlinking R-specific HTML files generated via pkgdown [12] directly to the Sphinx-generated documentation for Python and JupyterLab notebooks (as detailed in Methods B.1.4)—anticipating the growing integration of Python and R in bioinformatics.

By automating this documentation infrastructure, BasalCell ensures that every project built from its template is inherently “documentation-ready,” fostering greater reproducibility in biological research.

## BasalCell enables a seamless interoperable analysis pipeline

As a real-world example use case of BasalCell, we performed single-cell RNA-sequencing (scRNA-seq) data analysis, and deposited the full project directory to GitHub (https://github.com/yo-aka-gene/BasalCellDemo) for reference. As detailed in Supplementary Information (SI) A and Figure S1, BasalCell successfully orchestrated the Python and R environment to assure the interoperability between Python (e.g., `Scanpy` [13] and `CellTypist` [14]) and R (e.g., `clusterProfiler` [15] and `ComplexHeatmap` [16]) packages. By contrasting automated annotation leveraging Python’s machine-learning dominance with manual interpretation rooted in the comprehensive Bioconductor foundations in R, we demonstrate the inherent complementarity of these two paradigms and the potential for a true multi-language symbiosis in bioinformatics enabled by BasalCell.

## Discussion

A major advantage of BasalCell’s Jupyter-centric architecture is its inherent multi-language extensibility. Beyond Python and R, emerging high-performance languages such as Julia [17] and Mojo [18] are readily accessible via Jupyter kernels [19, 20]. In bioinformatics, leveraging these languages within a unified Jupyter environment holds great promise for computationally demanding tasks, including differential equation modeling, advanced mathematical optimization, and high-speed iterative processing.

By enhancing the robustness and reproducibility of biological data science, we envision BasalCell serving as a foundational scaffold to foster diverse biological discoveries in the future.

