## Supplemental Information for "BasalCell: A project scaffold generator for bioinformatics analysis"

### **Supplementary Information (SI)**: BasalCell: A project scaffold generator for bioinformatics analysis

#### Table of Contents

**A scRNA-seq data analysis with BasalCell** **ii**

**B Methods** **iv**

B.1 BasalCell implementation . . . . . iv

B.1.1 Cookiecutter template . . . . . iv

B.1.2 Virtual environment . . . . . iv

B.1.3 Continuous integration/continuous deployment (CI/CD) and code quality assurance . . . . . v

B.1.4 Documentation building . . . . . v

B.2 The peripheral blood mononuclear cell (PBMC) data analysis . . . . . vi

B.2.1 Quality control (QC) and preprocessing . . . . . vi

B.2.2 Differential expression analysis (DEA) . . . . . vi

B.2.3 Automated annotation with CellTypist . . . . . vi

B.2.4 Gene ontology (GO) analysis and manual annotation . . . . . vi

B.2.5 Heatmap visualization . . . . . vi

B.2.6 Data imput/output (I/O) between Python and R . . . . . vi

B.3 BasalCellBlank . . . . . vii

**C Data and code availability** **vii**

**D Author information** **vii**

#### A scRNA-seq data analysis with BasalCell

To demonstrate the seamless dual-language workflow, we created a project directory (named `BasalCellDemo`) and performed single-cell RNA-sequencing (scRNA-seq) data analysis. We used a publicly available scRNA-seq dataset of peripheral blood mononuclear cells (PBMCs). To highlight the benefits of integrating the computational efficiency of the Python ecosystem with the broad, specialized repertoire of the Bioconductor foundations in R, we compared the annotation labels derived from an automated approach using `CellTypist` [DCXJ+22] with a manual approach based on gene ontology (GO) enrichment analysis using `clusterProfiler` [YWHH12].

After preprocessing with `Scanpy` [WAT18], the PBMC cells were classified into seven clusters assigned by the Leiden algorithm [TWVE19] (Figure S1a). We then performed automated cell-type annotation using the `Immune_All_Low` model implemented in CellTypist (Figure S1b). To bridge the gap between the single-cell-level predictions provided by CellTypist and the cluster-wise manual annotations, we tallied the proportions of cell labels per cluster and assigned consensus identities based on the majority label within each cluster (Figures S1c–S1d).

For the manual annotation, we performed GO analysis using clusterProfiler following differential expression analysis (DEA) with Scanpy, and inferred the cell identities from the significantly enriched GO terms (Figure S1e). Notably, while we identified cluster 3 as cytotoxic T cells, we designated cluster 6 as undefined cellular identity due to ambiguous enrichment signatures (e.g., presentation of cytotoxicity and phagocytosis signatures).

To examine the discrepancies between the automated and manual annotations, we visualized the expression levels of the top 10 differentialy expressed geness (DEGs) for each cluster alongside the corresponding annotation labels using `ComplexHeatmap` (Figure S1f). Although the two annotation strategies achieved agreement across the majority of clusters, they exhibited distinctive classifications for clusters 3 and 6. Interestingly, a closer biological inspection reveals that both approaches effectively corrected each other's blind spots.

For cluster 3, while we observed high expression of genes indicative of cytotoxic T cells (e.g., *CD3D*, *CD3E*, *NKG7*, and *GZMA* [dlHMO+91, ZTB+17, TNT+25]), canonical mucosal-associated invariant T (MAIT)-specific markers (e.g., *KLRB1*, *SLC4A10*, *TRAV1-2*, and *DPP4* [DMS+11, HHAN+21, PK20]) were notably absent from the top 10 DEGs. Furthermore, although CellTypist predicted MAIT cells as the predominant type within this cluster, this assignment was highly competitive with other cell lineages (Figures S1e–S1f). Consequently, applying the more comprehensive "cytotoxic T cell" label via manual annotation is biologically more appropriate than adopting the automated MAIT cell classification.

Conversely, in cluster 6, alongside T cell-like cytotoxic markers (e.g., *NKG7*, *GNLY*, *GZMB*, and *PRF1* [ZTB+17, TNT+25]), signaling adaptors such as *TYROBP* and *FCER1G* were highly expressed alongside monocytic lineages. Because these adaptors mediate natural killer (NK) cell activation in addition to monocytic phagocytosis [HNK+09, LB00, ZWHL26], the interpretative difficulty of the GO terms in cluster 6 is elegantly resolved by the automated model's classification as NK cells.

Ultimately, this series of analyses highlights the mutual complementarity of the two ecosystems. The Python-based workflow, powered by modern machine learning models like CellTypist, enables rapid, data-driven annotation that effectively bypasses the bottleneck of manual human interpretation of DEGs. Conversely, the R-based workflow leverages the historical depth of the Bioconductor heritage; tools like clusterProfiler provide access to comprehensive semantic databases essential for nuanced biological inference, while packages like ComplexHeatmap offer the highly tailored visualizations indispensable for bioinformatics. By harmonizing these two distinctive milieus—computational agility and deeply contextualized, granular versatility—within a shared analytical workspace, BasalCell heralds the future of bioinformatics: a true multi-language symbiosis.

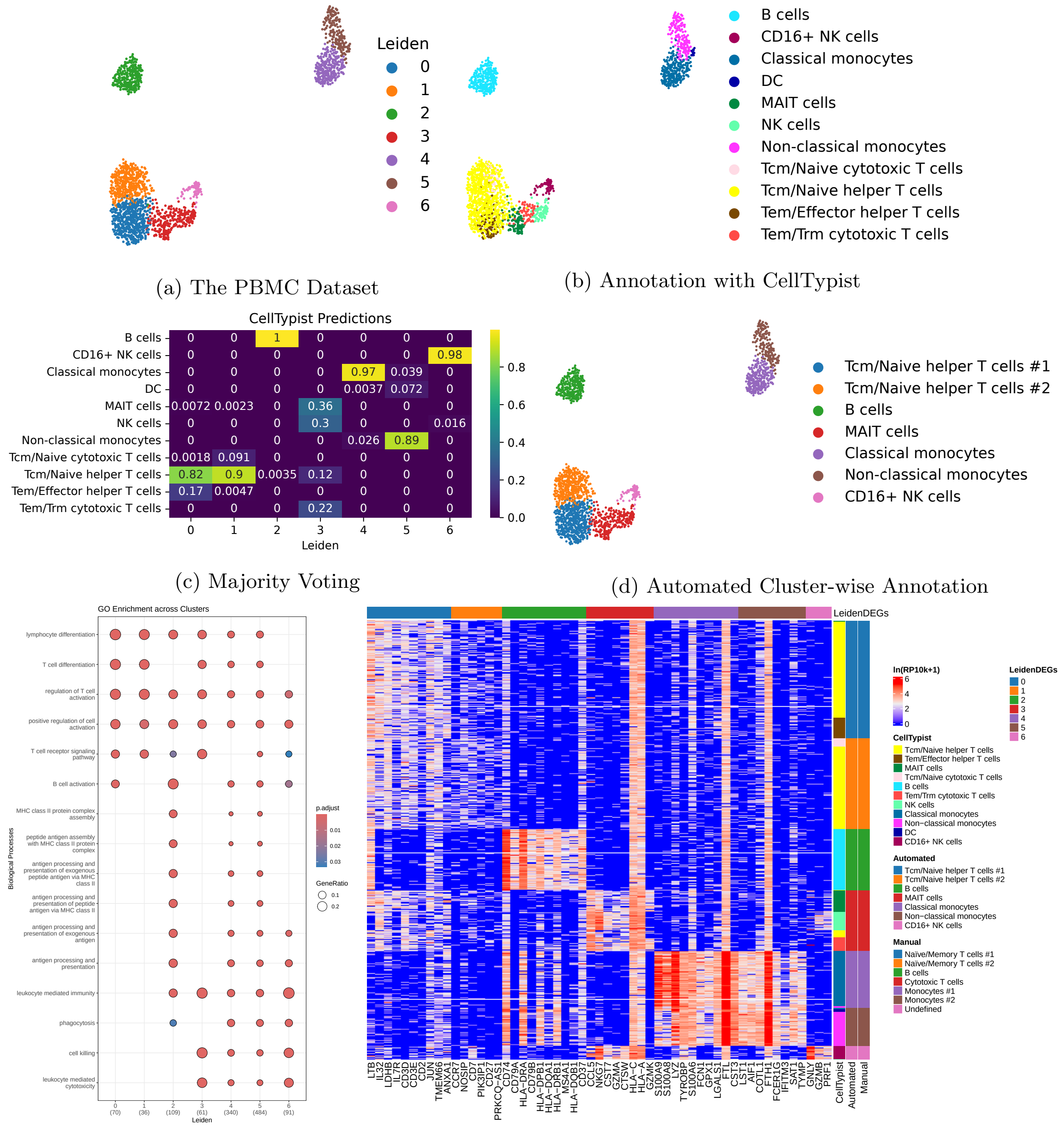


(a) The PBMC Dataset

(b) Annotation with CellTypist

(c) Majority Voting

(d) Automated Cluster-wise Annotation

(e) GO Analysis

(f) Heatmap created with ComplexHeatmap

**Figure S1: Comparative scRNA-seq annotation workflow utilizing Python and R ecosystems**

**a**: Uniform Manifold Approximation and Projection (UMAP) visualization of the PBMC dataset, identifying seven distinct clusters via the Leiden algorithm. **b**: Single-cell-level cell-type predictions generated by the automated CellTypist model. **c**: Proportional composition of the predicted cell labels within each Leiden cluster. **d**: Consensus automated annotations assigned to each cluster based on the majority label. **e**: Dot plot illustrating significantly enriched GO Biological Process terms for each cluster, computed using clusterProfiler. **f**: A comprehensive heatmap generated via ComplexHeatmap, displaying the expression profiles of the top 10 DEGs per cluster alongside the juxtaposition of the automated and manual annotation strategies. For visual clarity, overlapping genes among the top 10 DEGs across clusters were excluded from the cluster with the higher ID to eliminate redundancy.

### B Methods

#### B.1 BasalCell implementation

In this section, we outline the implementation procedures and underlying rationale for BasalCell and the BasalCellDemo repository. For comprehensive details regarding specific package versions, please refer to the `pyproject.toml` file for Python and the `renv.lock` file for R, both of which are available in their respective repositories (URLs are consolidated in Supplementary Information (SI) C).

##### B.1.1 Cookiecutter template

To achieve a disposable yet fully reproducible full-stack environment—encompassing version control, unit testing, linting, and automated documentation—we utilized the `cookiecutter` [RGc13] scaffold generator. While the technical specifics of each component are detailed in subsequent sections, the underlying template is driven by `Jinja` [Pal] templating syntax. This enables dynamic, prompt-based project generation, allowing users to flexibly tailor the environment to their specific needs.

Recognizing the ubiquity of Jupyter Lab in data science, BasalCell mandates Python as the primary analytical language. However, it provides modular options to include an R environment and/or initialize a Python package development directory, resulting in roughly four primary architectural permutations. For the R environment, users can not only toggle its inclusion but also specify the language version (4.3 or 4.4), ensuring precise configuration for R-based analytical pipelines.

Regarding Python, BasalCell currently enforces version 3.12 exclusively. This restriction stems from distinct compatibility challenges associated with versions $\leq 3.11$ and $\geq 3.13$.

First, regarding Python 3.11 and earlier, significant version conflicts arise within foundational packages. BasalCell is designed so that researchers without deep software engineering expertise can easily construct their environments; thus, the expected default use case is that users will install packages without explicitly pinning versions. Under these conditions, `Poetry` [ET] attempts to resolve dependencies and install the latest compatible versions. However, critical low-level dependencies like `Numba` [Numb] and `llvmlite` [Numa]—which are fundamental to core bioinformatics libraries such as `NumPy` [HMVDW$^+$20], `pandas` [M$^+$10], `SciPy` [VGO$^+$20], `Scanpy` [WAT18], and `Polars` [Vin24]—exhibit divergent compatibility profiles between Python $\leq 3.11$ and $\geq 3.12$. Consequently, Poetry's dependency solver often proposes complex solutions that attempt to force packages requiring newer `Numba`/`llvmlite` versions into a legacy Python 3.11 environment. This inevitably leads to installation failures, forcing the user to manually intervene and reconstruct the entire dependency tree. Requiring such advanced troubleshooting contradicts BasalCell's core mission of democratizing rigorous development environments, prompting us to deprecate support for Python $\leq 3.11$.

Second, regarding Python 3.13 and newer, our chosen high-performance linter, `Ruff` [Mc], currently lacks full stable support for these latest releases. To maintain the robustness of the CI/CD pipeline, we deferred 3.13 integration.

Although Python 3.12 is the sole supported version in the current release (`v0.4.1`) due to these constraints, the interactive cookiecutter prompt retains a version selection menu. This design choice ensures structural forward-compatibility, allowing seamless integration once the surrounding toolchain fully adapts to future Python releases.

##### B.1.2 Virtual environment

To ensure code reproducibility, we identified three essential pillars: (i) establishing isolated development environments for each project, (ii) maintaining full transparency of software dependency versions, and (iii) guaranteeing the overall reproducibility and distributability of the workspace.

Based on these principles, we adopted a virtual environment architecture built on `Mamba` [QM19]. Using cookiecutter, user-specified versions of the Python and R languages are pinned within a virtual environment initialized via `Miniforge` [con20]. Within this environment, `Poetry` [ET] manages Python package dependencies independently while coordinating with Mamba to ensure language-level consistency.

For the R environment, although `renv` [UW26] is a standard for version management, it often encounters performance bottlenecks and conflicts with the host system due to complex OS-level dependencies and compilation requirements. To mitigate these issues, we implemented a hybrid management strategy: primarily, pre-compiled R packages and their associated OS-level dependencies are installed via Mamba, while renv is utilized strictly as a tracking mechanism to record precise package versions. This approach significantly accelerates environment restoration and minimizes compilation failures. For edge cases where specific versions are unavailable on Conda channels or for packages under active development (e.g., hosted on GitHub), the system falls back to direct installation via renv, ensuring both stability and flexibility. Consequently, the complete blueprint for R packages is captured by `renv.lock`, whereas OS-level dependencies and the overarching environment definitions are maintained by `conda-lock.yml`.

However, maintaining independent lock files for Mamba and renv inherently risks version management collisions. To resolve this, BasalCell parses the configuration files of Mamba, Poetry, and renv to construct a unified dataframe. By programmatically ensuring Mamba always takes precedence over renv during operations, the system avoids package manager conflicts. This design successfully harmonizes Mamba's rapid installation and OS-level stability with the rigorous version tracking of renv. Furthermore, this integrated dataframe is constructed using `Polars` [Vin24], enabling high-speed querying and versatile data export into multiple formats (CSV, IPC, Parquet). We also implemented a feature to export queried version data directly as Markdown files, significantly enhancing human readability and methodological accessibility of static configuration data.

Regarding OS-level dependencies, while strict binary-level locking is ideal from the perspective of code reproducibility, such an approach often hinders cross-platform portability across different operating systems. Therefore, to maximize reproducibility while ensuring cross-platform compatibility, the `conda-lock.yml` file is configured to simultaneously lock versions across three major architectures: `linux-64`, `osx-arm64`, and `osx-64`.

Finally, to simplify user operations, complex multi-line command sequences—encompassing environment initialization and unified workflows for software installation and version logging—are encapsulated using `GNU Make` [SMS98].

This multi-layered architecture achieves comprehensive reproducibility across the Python, R, and OS tiers without sacrificing workspace portability.

##### B.1.3 Continuous integration/continuous deployment (CI/CD) and code quality assurance

To establish a robust CI/CD pipeline, BasalCell implements automated linting and code testing both in the local development environment and via GitHub Actions. This dual-layered approach ensures consistent code quality and readability for both individual and collaborative projects.

For linting, we integrated `Ruff` [Mc] for Python, alongside `lintr` [HAC+25] and `styler` [MWP26] for R. By orchestrating these tools through `pre-commit` [SKc] hooks, BasalCell automatically triggers code formatting and linting upon every Git commit. This design intentionally enforces standard software engineering practices implicitly, empowering even users unfamiliar with these concepts to maintain clean codebases. Furthermore, GitHub Actions provides a secondary validation layer by automatically executing this same linting workflow upon any push or pull request to the main branch.

In contrast, BasalCell adopts a highly flexible, non-restrictive approach for code testing. In bioinformatics, coding practices span a wide spectrum—from fluid exploratory data analysis to robust package development—meaning the required level of code rigor (e.g., strict typing or handling complex edge cases) varies significantly. While test-driven development (TDD) is a cornerstone of software engineering, enforcing production-level testing for all biological coding tasks imposes an unnecessary and impractical burden on researchers. Specifically, decomposing analytical processes into isolated components, anticipating all conceivable valid and invalid inputs, and writing exhaustive unit tests—to validate that the implementation behaves exactly as the requirements definition specifies under every condition—presents a formidable barrier for those unfamiliar with TDD. Furthermore, even for experienced developers, this rigorous testing demands substantial time and labor. Consequently, whether such strict unit testing is truly justified from a cost-benefit perspective depends entirely on the specific context and lifespan of the analytical code.

Therefore, local code testing in BasalCell is entirely optional. When testing is desired, the framework natively supports lightweight function validation via Python's `doctest`. Simultaneously, it fully supports rigorous unit testing via `pytest` [KOP+04] and `testthat` [Wic11], accommodating scenarios demanding strict code reliability, such as package development. However, for collaborative development, GitHub Actions enforces automated code testing (running both doctests and formal unit tests) upon any push or pull request to the main branch, acting as a strict quality control checkpoint. Notably, if no tests are implemented, the GitHub Actions workflow is designed to fail, visibly indicating on the repository that the codebase has not passed a testing pipeline.

##### B.1.4 Documentation building

To enhance methodological transparency, we prioritized code readability as a fundamental requirement for peer review and reproducible research. We identified three core benefits of documentation for non-package bioinformatics projects: (i) clarifying analytical intent and context, (ii) organizing disparate scripts into a logical narrative, and (iii) restructuring idiosyncratic directory architectures into intuitive web-based interfaces.

To lower the implementation barrier, BasalCell implements an automated documentation workflow using `Sphinx` [KTc] and Read the Docs. We adopted `JupyterLab` [KRKP+16] as the primary analytical interface due to its native support for Markdown-based annotations, strict correspondence between code and output, and multi-language kernel extensibility. Crucially, BasalCell adopts a deployment strategy wherein locally generated HTML artifacts are uploaded directly to the Read the Docs server, deliberately bypassing server-side re-rendering. By explicitly configuring the build engine to skip notebook execution, the exact computational outputs and experimental logs generated in the user's local environment are statically preserved. This ensures that JupyterLab notebook files function directly as immutable documentation, significantly enhancing the traceability and readability of the analytical workflow.

While Sphinx is the standard for Python, it lacks native support for R script documentation. To bridge this gap, BasalCell utilizes `pkgdown` [WHS18] to generate HTML documentation for R scripts, which are then integrated into the Sphinx ecosystem via hyperlinking within the Markdown-based index. Furthermore, the framework supports standard R-package vignettes using `rmarkdown` [AXD+26] and `knitr` [Xie25]. To further improve script-level interpretability, BasalCell leverages `docstrings` for Python and `roxygen2` [WDCE25] for R, both of which are automatically rendered into HTML via their respective documentation engines.

#### B.2 The PBMC data analysis

##### B.2.1 Quality control (QC) and preprocessing

The PBMC dataset was obtained from 10x Genomics public sources (see SI C for details). Using `Scanpy` [WAT18], we performed QC and filtered cells based on multi-parametric criteria, including the number of genes per cell, total counts, and the percentages of mitochondrial, ribosomal, and hemoglobin genes, implemented via `Polars` [Vin24]. After doublet removal using `Scrublet` [WLK19] within Scanpy, raw counts were normalized using a $\ln(\mathrm{RP10k} + 1)$ transformation.

Dimensionality reduction was performed using principal component analysis (PCA) (first 50 principal components) on the top 3,000 highly variable geness (HVGs). Subsequently, a neighborhood graph was constructed, followed by Uniform Manifold Approximation and Projection (UMAP) embedding and Leiden clustering.

##### B.2.2 DEA

DEA was conducted using the Wilcoxon rank-sum test based on the clustering results described in Methods B.2.1.

##### B.2.3 Automated annotation with CellTypist

Automated cell-type annotation was performed at the single-cell level using the `Immune_All_Low` model in CellTypist [DCXJ+22]. For cluster-wise annotation, predictions were aggregated per cluster, and each cluster was assigned the identity of its most frequent cell type (majority voting).

##### B.2.4 GO analysis and manual annotation

Following DEA, genes with adjusted $p$-values $< 0.05$ and $\log_2(\mathrm{RPM}{+}1) > 1$ were extracted using Polars. GO enrichment analysis was performed for each cluster using clusterProfiler [YWHH12] with $p$-value and $q$-value thresholds set to 0.05. Manual annotations were subsequently inferred based on the significantly upregulated GO terms.

##### B.2.5 Heatmap visualization

The expression levels of the top 10 DEGs for each cluster, along with the results of automated and manual annotations, were visualized using ComplexHeatmap [GES16]. For visual clarity, overlapping genes among the top 10 DEGs across clusters were excluded from the cluster with the higher ID to eliminate redundancy.

##### B.2.6 Data imput/output (I/O) between Python and R

To achieve high-speed and memory-efficient data I/O between Python and R, relevant analytical information from Scanpy and Polars was converted into data frames and exported as Parquet files using Polars, preserving strict type information. These files were subsequently retrieved in R using the `arrow` [RCC+24] package.

##### B.3 BasalCellBlank

To demonstrate the unmodified output of BasalCell, we generated a baseline project directory. The generation process was performed by selecting the following options in the interactive CLI:

**Project name:** BasalCellBlank

**Description:** Just Generated via BasalCell

**Author:** Yuji Okano

**GitHub username:** yo-aka-gene

**Python version:** 3.12

**R version:** 4.4 (Option 3)

**Package structure:** Enabled (Option 2: true)

The resulting directory was deposited to GitHub (https://github.com/yo-aka-gene/BasalCellBlank) to serve as a reference for the generator's default state without any manual modifications.

#### C Data and code availability

The PBMC scRNA-seq dataset is publicly available from the online resource (https://support.10xgenomics.com/single-cell-gene-expression/datasets/1.1.0/pbmc3k).

The source code of BasalCell, the scRNA-seq data analysis code and the environment configurations in BasalCellDemo, and the original state of a project directory generated with BasalCell (`BasalCellBlank`) are all deposited in GitHub:

- BasalCell: https://github.com/yo-aka-gene/BasalCell
- BasalCellDemo: https://github.com/yo-aka-gene/BasalCellDemo
- BasalCellBlank: https://github.com/yo-aka-gene/BasalCellBlank

#### D Author information

YO conceived the method, implemented the package and analysis code with input from TI. YO and TI performed debugging. YO wrote the manuscript with input from TI and KS. All the authors reviewed the manuscript.

#### Supplemental References


[AXD+26] JJ Allaire, Yihui Xie, Christophe Dervieux, Jonathan McPherson, Javier Luraschi, Kevin Ushey, Aron Atkins, Hadley Wickham, Joe Cheng, Winston Chang, and Richard Iannone. *rmarkdown: Dynamic Documents for R*, 2026. R package version 2.31.

[con20] conda-forge community. Miniforge. https://github.com/conda-forge/miniforge, 2020. GitHub repository.

[DCXJ+22] César Domínguez Conde, Chao Xu, Louie B Jarvis, Daniel B Rainbow, Sara B Wells, Tamir Gomes, SK Howlett, O Suchanek, K Polanski, HW King, et al. Cross-tissue immune cell analysis reveals tissue-specific features in humans. *Science*, 376(6594):eabl5197, 2022.

[dlHMO+91] Antonio de la Hera, U Müller, Carina Olsson, Sylvie Isaaz, and Alan Tunnacliffe. Structure of the T cell antigen receptor (TCR): two CD3 epsilon subunits in a functional TCR/CD3 complex. *The Journal of experimental medicine*, 173(1):7–17, 1991.

[DMS+11] Mathilde Dusseaux, Emmanuel Martin, Nacer Serriari, Isabelle Péguillet, Virginie Premel, Delphine Louis, Maud Milder, Lionel Le Bourhis, Claire Soudais, Emmanuel Treiner, et al. Human MAIT cells are xenobiotic-resistant, tissue-targeted, CD161hi IL-17–secreting T cells. *Blood, The Journal of the American Society of Hematology*, 117(4):1250–1259, 2011.

[ET] Sébastien Eustace and The Poetry contributors. Poetry: Python packaging and dependency management made easy. https://python-poetry.org. GitHub repository: https://github.com/python-poetry/poetry.

[GES16] Zuguang Gu, Roland Eils, and Matthias Schlesner. Complex heatmaps reveal patterns and correlations in multidimensional genomic data. *Bioinformatics*, 32(18):2847–2849, 2016.

[HAC+25] Jim Hester, Florent Angly, Michael Chirico, Russ Hyde, Ren Kun, Indrajeet Patil, and Alexander Rosenstock. Static Code Analysis for R. *Journal of Open Source Software*, 10(108):7240, 2025.

[HHAN+21] Yuhan Hao, Stephanie Hao, Erica Andersen-Nissen, William M Mauck, Shiwei Zheng, Andrew Butler, Maddie J Lee, Aaron J Wilk, Charlotte Darby, Michael Zager, et al. Integrated analysis of multimodal single-cell data. *Cell*, 184(13):3573–3587, 2021.

[HMVDW+20] Charles R Harris, K Jarrod Millman, Stéfan J Van Der Walt, Ralf Gommers, Pauli Virtanen, David Cournapeau, Eric Wieser, Julian Taylor, Sebastian Berg, Nathaniel J Smith, et al. Array programming with NumPy. *Nature*, 585(7825):357–362, 2020.

[HNK+09] Jessica A Hamerman, Minjian Ni, Justin R Killebrew, Ching-Liang Chu, and Clifford A Lowell. The expanding roles of ITAM adapters FcR$\gamma$ and DAP12 in myeloid cells. *Immunological reviews*, 232(1):42–58, 2009.

[KOP+04] Holger Krekel, Bruno Oliveira, Ronny Pfannschmidt, Floris Bruynooghe, Brianna Laugher, Freya Bruhin, and contributors. pytest. https://github.com/pytest-dev/pytest, 2004. GitHub repository.

[KRKP+16] Thomas Kluyver, Benjamin RAGAN-KELLEYb[1], Fernando Pérez, Brian Granger, Matthias Bussonnier, Jonathan Frederic, and Kyle Kelley. Jupyter Notebooks–a publishing format for reproducible computational workflows. In *Positioning and power in academic publishing: players, agents and agendas: proceedings of the 20th International Conference on Electronic Publishing*, page 87. IOS press, 2016.

[KTc] Takeshi Komiya, Adam Turner, and contributors. Sphinx: The Sphinx documentation generator. https://github.com/sphinx-doc/sphinx. GitHub repository.

[LB00] Lewis L Lanier and Alexander BH Bakker. The ITAM-bearing transmembrane adaptor DAP12 in lymphoid and myeloid cell function. *Immunology today*, 21(12):611–614, 2000.

[M+10] Wes McKinney et al. Data structures for statistical computing in python. In *Proceedings of the 9th Python in Science Conference*, volume 445, pages 51–56. Austin, TX, 2010.

[Mc] Charlie Marsh and contributors. Ruff: An extremely fast Python linter and code formatter, written in Rust. https://github.com/astral-sh/ruff. GitHub repository.

[MWP26] Kirill Müller, Lorenz Walthert, and Indrajeet Patil. *styler: Non-Invasive Pretty Printing of R Code*, 2026. R package version 1.11.0.

[Numa] Numba Developers. llvmlite: A lightweight LLVM python binding for writing JIT compilers. https://github.com/numba/llvmlite. GitHub repository.

[Numb] Numba Developers. Numba: NumPy aware dynamic Python compiler using LLVM. https://github.com/numba/numba. GitHub repository.

[Pal] Pallets Projects. Jinja: A very fast and expressive template engine. https://github.com/pallets/jinja. GitHub repository.

[PK20] Nicholas M Provine and Paul Klenerman. Mait cells in health and disease. *Annual review of immunology*, 38(1):203–228, 2020.

[QM19] QuantStack and Mamba Contributors. Mamba: The Fast Cross-Platform Package Manager. https://github.com/mamba-org/mamba, 2019. GitHub repository.

[RCC+24] Neal Richardson, Ian Cook, Nic Crane, Dewey Dunnington, Romain François, Jonathan Keane, Dragoș Moldovan-Grünfeld, Jeroen Ooms, Jacob Wujciak-Jens, and Apache Arrow. *arrow: Integration to 'Apache' 'Arrow'*, 2024. R package version 15.0.0, https://arrow.apache.org/docs/r/.

[RGc13] Audrey M. Roy Greenfeld and contributors. Cookiecutter. https://github.com/cookiecutter/cookiecutter, 2013. GitHub repository.

[SKc] Ken Struys, Chris Kuehl, and contributors. pre-commit: A framework for managing and maintaining multi-language pre-commit hooks. https://github.com/pre-commit/pre-commit. GitHub repository.

[SMS98] Richard M. Stallman, Roland McGrath, and Paul D. Smith. *GNU Make*. Free Software Foundation, 1998.

[TNT+25] Roberta Turiello, Susanna S Ng, Elisabeth Tan, Gemma van der Voort, Nazhifah Salim, Michelle CR Yong, Malika Khassenova, Johannes Oldenburg, Heiko Rühl, Jan Hasenauer, et al. NKG7 is a stable marker of cytotoxicity across immune contexts and within the tumor microenvironment. *European Journal of Immunology*, 55(6):e51885, 2025.

[TWVE19] Vincent A Traag, Ludo Waltman, and Nees Jan Van Eck. From louvain to leiden: guaranteeing well-connected communities. *Scientific reports*, 9(1):5233, 2019.

[UW26] Kevin Ushey and Hadley Wickham. *renv: Project Environments*, 2026. R package version 1.2.1.

[VGO+20] Pauli Virtanen, Ralf Gommers, Travis E Oliphant, Matt Haberland, Tyler Reddy, David Cournapeau, Evgeni Burovski, Pearu Peterson, Warren Weckesser, Jonathan Bright, et al. SciPy 1.0: fundamental algorithms for scientific computing in Python. *Nature methods*, 17(3):261–272, 2020.

[Vin24] Ritchie Vink. Polars: Extremely fast Query Engine for DataFrames, written in Rust. https://github.com/pola-rs/polars, 2024. GitHub repository.

[WAT18] F Alexander Wolf, Philipp Angerer, and Fabian J Theis. SCANPY: large-scale single-cell gene expression data analysis. *Genome biology*, 19(1):15, 2018.

[WDCE25] Hadley Wickham, Peter Danenberg, Gábor Csárdi, and Manuel Eugster. *roxygen2: In-Line Documentation for R*, 2025. R package version 7.3.3.

[WHS18] Hadley Wickham, Jay Hesselberth, and Maëlle Salmon. Pkgdown: Make static html documentation for a package. *R package version*, 1(0), 2018.

[Wic11] Hadley Wickham. testthat: Get Started with Testing. *The R Journal*, 3:5–10, 2011.

[WLK19] Samuel L Wolock, Romain Lopez, and Allon M Klein. Scrublet: computational identification of cell doublets in single-cell transcriptomic data. *Cell systems*, 8(4):281–291, 2019.

[Xie25] Yihui Xie. *knitr: A General-Purpose Package for Dynamic Report Generation in R*, 2025. R package version 1.51.

[YWHH12] Guangchuang Yu, Li-Gen Wang, Yanyan Han, and Qing-Yu He. clusterProfiler: an R package for comparing biological themes among gene clusters. *Omics: a journal of integrative biology*, 16(5):284–287, 2012.

[ZTB+17] Grace XY Zheng, Jessica M Terry, Phillip Belgrader, Paul Ryvkin, Zachary W Bent, Ryan Wilson, Solongo B Ziraldo, Tobias D Wheeler, Geoff P McDermott, Junjie Zhu, et al. Massively parallel digital transcriptional profiling of single cells. *Nature communications*, 8(1):14049, 2017.

[ZWHL26] Yuyu Zhang, Jinglan Wang, Wenting He, and Tao Liu. FCER1G: A multifunctional regulator in the immune microenvironment. *Experimental and Therapeutic Medicine*, 31(5):146, 2026.
